# Robust RNA editing via recruitment of endogenous ADARs using circular guide RNAs

**DOI:** 10.1101/2021.01.12.426286

**Authors:** Dhruva Katrekar, James Yen, Yichen Xiang, Anushka Saha, Dario Meluzzi, Yiannis Savva, Prashant Mali

## Abstract

Akin to short-hairpin RNAs and antisense oligonucleotides which efficaciously recruit endogenous cellular machinery such as Argonaute and RNase H to enable targeted RNA knockdown, simple long antisense guide RNAs (*1*) can recruit endogenous adenosine deaminases acting on RNA (ADARs) to enable programmable A-to-I RNA editing, without requiring co-delivery of any exogenous proteins. This approach is highly specific, however the efficiency is typically lower than observed with enzyme overexpression. Conjecturing this was due in part to the short half-life and residence times of guide RNAs, here we engineer highly stable <u>c</u>ircular <u>AD</u>AR recruiting guide <u>RNAs</u> (cadRNAs), which can be delivered not only by genetically encoding on DNA vectors, but also via transfection of RNA molecules transcribed *in vitro*. Using these cadRNAs, we observed robust RNA editing across multiple sites and cell lines, in both untranslated and coding regions of RNAs, vastly improved efficiency and durability of RNA editing, and high transcriptome-wide specificity. High transcript-level specificity was achieved by further engineering to reduce bystander editing. Additionally, *in vivo* delivery of cadRNAs via adeno-associated viruses (AAVs) enabled robust 38% RNA editing of the mPCSK9 transcript in C57BL/6J mice livers, and 12% UAG-to-UGG RNA correction of the amber nonsense mutation in the IDUA-W392X mouse model of mucopolysaccharidosis type I-Hurler (MPS I-H) syndrome. Taken together, cadRNAs enable efficacious programmable RNA editing with application across diverse protein modulation and gene therapeutic settings.

## INTRODUCTION

Adenosine to inosine (A-to-I) RNA editing is a common post-transcriptional modification catalyzed by adenosine deaminases acting on RNA (ADAR) enzymes (*2*–*8*). ADARs edit double stranded RNA (dsRNA) predominantly in non-coding regions such as Alu repetitive elements in a promiscuous fashion, while also editing a handful of sites in coding regions with high specificity (*9*–*13*). The structural similarity between inosine and guanosine results in the translation and splicing machinery recognizing the edited base as guanosine, thereby making ADARs attractive tools for recoding protein sequences (*14*). To this end, several studies have recently repurposed the ADAR system for programmable RNA editing both *in vitro* (*1, 15–22*) and *in vivo* (*1, 23*) by engineering recruitment of ADARs to a target RNA sequence using ADAR recruiting guide RNAs (adRNAs). Although ADARs, and in particular ADAR1, are widely expressed throughout the body, most of these studies relied on exogenously delivered ADAR enzymes and their variants to achieve robust RNA editing efficiencies. However, as ADAR-dsRNA interactions primarily rely on structure rather than sequence dependency, a major limitation of relying on enzyme overexpression is the propensity to introduce a plethora of off-target A-to-I edits across the transcriptome (*1, 19, 24, 25*). Additionally, as ADARs are native to and thus not orthogonal to most mammalian systems, their overexpression can result in altered protein interactions that might impact cellular physiology. Furthermore, as this approach relies on two components, a guide RNA and the ADAR protein, it can limit delivery modalities, in particular for *in vivo* applications.

A solution to this is to engineer adRNAs to enable recruitment of endogenous ADARs. Towards this, we recently demonstrated that it is possible to recruit endogenous ADARs using simple long antisense RNA of length >60 bp (*1*). This strategy is exciting since akin to short-hairpin RNAs (shRNAs) and antisense oligonucleotides (ASOs) which efficaciously recruit endogenous cellular machinery such as Argonaute (*26*) and RNase H (*27, 28*) to enable targeted RNA knockdown, just delivery of guide RNAs alone can now enable programmable A-to-I RNA editing without requiring co-delivery of any exogenous proteins. However, the efficiency of RNA editing via this approach is typically lower than seen with enzyme overexpression, thus limiting their utility in biotechnology and therapeutic applications. Conjecturing this was due in part to the short half-life and residence times of guide RNAs, here we engineer highly stable circular ADAR recruiting guide RNAs (cadRNAs). These vastly improve the efficiency and durability of RNA editing. We demonstrate too that targeting via cadRNAs is highly specific at the transcriptome-wide level, and via further engineering to reduce bystander editing, also highly specific at the transcript level. Furthermore, we show cadRNAs can be delivered genetically encoded via DNA, and as well via *in vitro* transcribed RNA at a fraction of the cost of chemically synthesized ASOs. Additionally, these enable highly robust RNA editing in both untranslated and coding regions of mRNAs, and across multiple RNA targets and cell lines. Importantly, using cadRNAs, we also demonstrate for the first time robust *in vivo* RNA editing via endogenous ADAR recruitment, including in the IDUA-W392X mouse model of mucopolysaccharidosis type I-Hurler (MPS I-H) syndrome.

## RESULTS

Using as the base format our long antisense guide RNA design that can recruit endogenous ADARs, we explored two guide RNA engineering strategies to further enhance RNA editing efficiencies (**Figure 1a**): one, we coupled recruiting domains that are derived from native RNAs sites known to be heavily edited by ADARs; and two, we coupled domains that stabilize and confer increased half-life of the guide RNAs.

**Figure 1:**
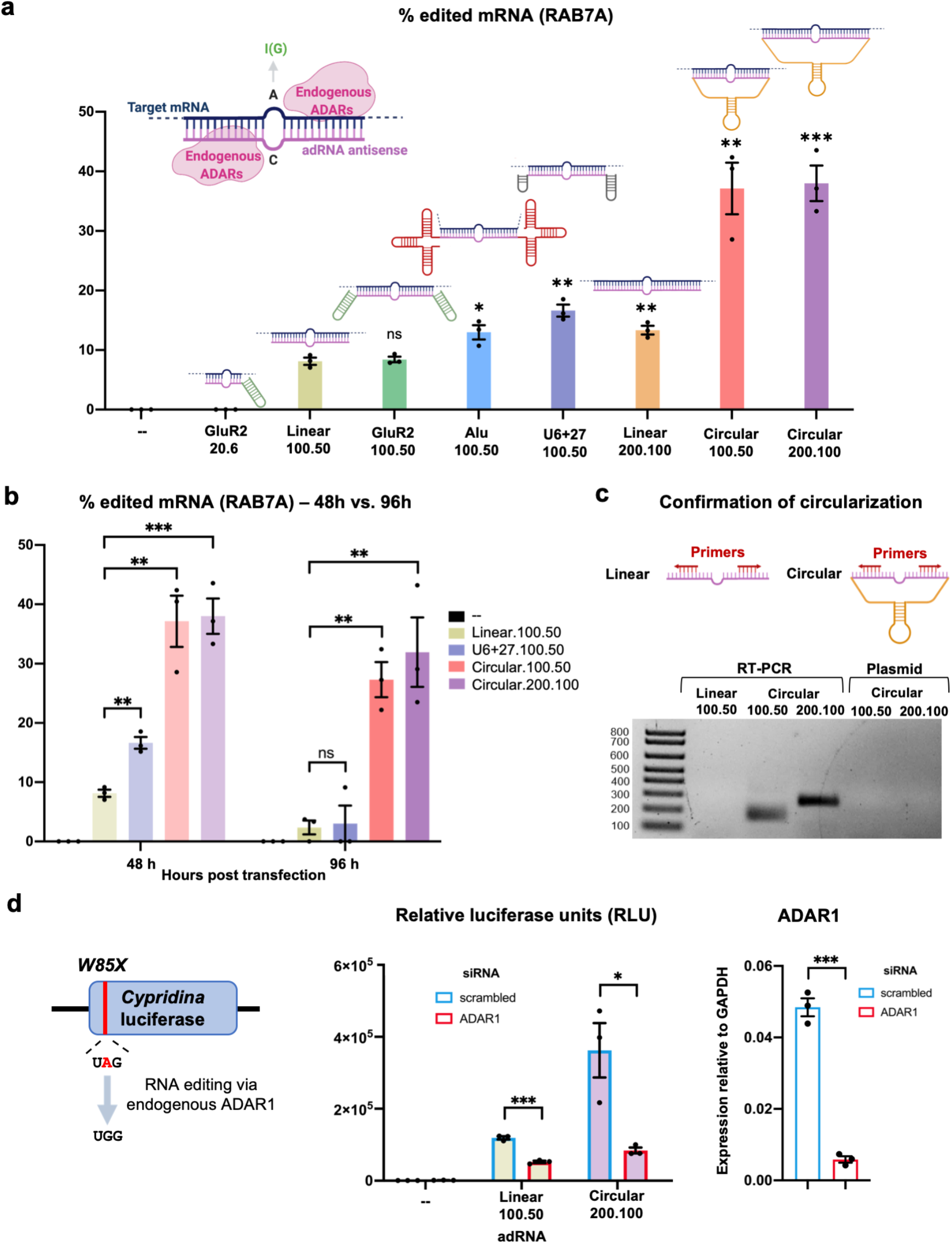
Engineering circular ADAR recruiting guide RNAs (cadRNAs). (**a**) A comparison of the RNA editing efficiencies in the 3’ UTR of the RAB7A transcript via various adRNA designs. Values represent mean +/- SEM (n=3; with respect to the linear.100.50, left-to-right, p=0.7289, p=0.0226, p=0.0019, p=0.0055, p=0.0027, and p=0.0006; unpaired t-test, two-tailed). In the schematics, the pink strand represents the antisense domain of the adRNA while the target mRNA is in blue. The bulge indicates the A-C mismatch between the target mRNA and adRNA. (**b**) RNA editing efficiencies achieved 48 hours and 96 hours post transfection of various adRNA designs. Values represent mean +/- SEM (n=3; left-to-right, p=0.0019, p=0.0027, p=0.0006 and p=0.8488, p=0.0014, p=0.0077; unpaired t-test, two-tailed). The 48 hour panel data is reproduced from Figure 1a>. (**c**) RT-PCR based confirmation of adRNA circularization in cells. (**d**) The ability of adRNAs to effect RNA editing of the cluc transcript was assessed in the presence of an siRNA targeting ADAR1. Values represent mean +/- SEM (n=3; left-to-right, p=0.0002, p=0.0216 and p=0.0001; unpaired t-test, two-tailed). All experiments were carried out in HEK293FT cells.

Towards the former we evaluated recruiting domains from the naturally occurring ADAR2 substrate GluR2 pre-messenger RNA (*17, 18*), and Alu elements which are known substrates for ADAR1 (*29*). The Alu adRNAs were created by positioning the antisense domain within the Alu consensus sequence. We screened these modified guide RNAs by assaying editing at an adenosine in the 3’UTR of the RAB7A transcript in HEK293FT cells. Consistent with our previous observations (*1*), the GluR2 domain coupled to a short antisense of length 20bp (GluR2.20.6) was unable to recruit endogenous ADARs resulting in no detectable RNA editing, while, as we have previously shown, the long antisense RNAs (linear.100.50) alone resulted in modest ∼10% RNA editing. Coupling the GluR2 domains to the long antisense version (GluR2.100.50) did not enhance RNA editing yields, but we observed that the addition of Alu domains (Alu.100.50) marginally enhanced the efficiency of RNA editing (1.5-fold). While significant, these designs had only a modest improvement over the simple long antisense versions.

We thus focused next on evaluating the impact of persistence of adRNAs, as this in turn could also impact target RNA search as well as their net target residence times. In particular, genetically encoded adRNAs are typically expressed via the polymerase III promoter, and thus transcribed guides lack a 5’ cap and a 3’ poly A tail and correspondingly have very short half-lives. To improve guide RNA persistence we evaluated: 1) increasing the length of the guide RNAs (linear.200.100); 2) coupling a U6+27 cassette (U6+27.100.50) which has been shown to improve stability of siRNA (*30*); and 3) engineering circularized versions (circular.100.50 and circular.200.100) as these would be intrinsically resistant to cellular exonucleases. Specifically, leveraging an elegant methodology recently developed by Litke and colleagues (*31*), we engineered circular ADAR recruiting guide RNAs (cadRNAs) by flanking the linear adRNAs by twister ribozymes, which upon autocatalytic cleavage leave termini that are ligated by the ubiquitous endogenous RNA ligase RtcB to yield circularized guide RNAs. Comparing the three different guide designs we observed that both increasing the adRNA length and the addition of U6+27 to the long antisense adRNA led to a 1.5-fold and 2-fold respective improvement in editing of the RAB7A transcript over the linear.100.50 designs (**Figure 1a**). Notably, using circular adRNA with antisense lengths 100bp and 200bp (i.e. circular.100.50 and circular.200.100), resulted in an even more robust 3.5-fold improvement in efficiency over the linear.100.50 designs and a 2-fold improvement over the Alu.100.50 and U6+27.100.50 designs (**Figure 1a**). Excitingly, we observed persistence of significant levels in RNA editing at both 48 hours and 96 hours post transfection, while editing via linear guide RNAs was almost non-detectable by the 96 hour time point (**Figure 1b**). We confirmed covalent circularization of the cadRNAs in HEK293FT cells via RT-PCR by designing outward facing primers that selectively amplified only the circularized structure (**Figure 1c**). Finally, we confirmed that RNA editing via the circular guide RNAs, similar to the linear guide RNAs, was mediated by endogenous ADAR1 recruitment. Towards this, we performed a luciferase based reporter assay, where we assayed the guide RNAs for their ability to repair a premature stop codon (UAG) in the *cypridina* luciferase (cluc) transcript (*19*) in the presence of scrambled and ADAR1 specific siRNAs. We observed a significant drop in luciferase activity in the presence of ADAR1 siRNA, confirming that RNA editing via long antisense adRNAs and circular adRNAs was dependent on endogenous ADAR1 levels (**Figure 1d**).

We next sought to evaluate the specificity profile of cadRNAs at both the transcriptome wide and target transcript levels. Towards the former, a circular.100.50 and a circular.200.100 sample along with an untransfected HEK293FT sample were analyzed by deep RNA-seq. Notably, in contrast with enzyme overexpression where we routinely observed 10^3^-10^4^ transcriptome wide off-targets (*1*), we noted 2-3 orders of magnitude lower off-target editing via the cadRNAs and at levels similar to the linear long antisense guide RNAs (**Figure 2a**). At the transcript level, however, we did observe bystander editing at flanking adenosines (**Figure 2b**). This is attributable to the long and perfect paired dsRNA stretch created upon adRNA-target binding. By creating a G mismatch (*32*) opposite all non-target adenosines we could completely eliminate this bystander editing, however this also led to a significant drop in the on-target editing efficiency to about 35% of the unmodified circular.200.100 version (**Figure 2b**). To address this, we engineered the antisense region to more closely mimic dsRNA structures of natural ADAR substrates. Specifically, we engineered 8bp bulge loops positioned both 6bp upstream and 30bp downstream of the target (*33*). This novel design led to a significant reduction in bystander editing, with the on-target editing being double that achieved by simply placing opposing G mismatches (**Figure 2b**). Taken together, this combination of 8bp loops to create breaks within the long stretch of dsRNA, combined with certain A-specific bulges can thus be utilized to reduce bystander editing.

**Figure 2:**
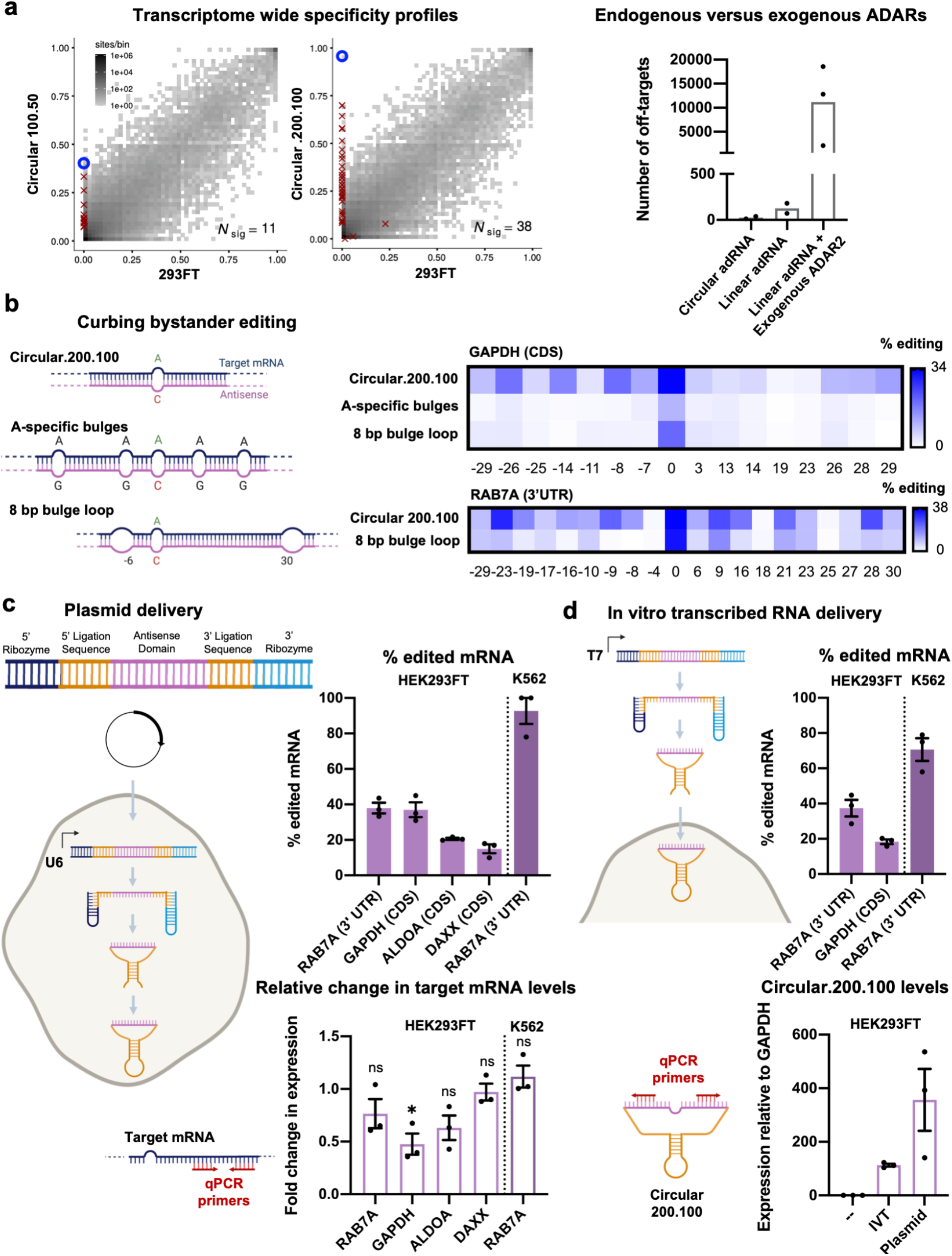
*In vitro* efficacy of cadRNAs. (**a**) (left-panel) 2D histograms comparing the transcriptome-wide A-to-G editing yields observed with a circular adRNA construct (*y*-axis) to the yields observed with the control sample (*x*-axis). Each histogram represents the same set of reference sites, where read coverage was at least 10 and at least one putative editing event was detected in at least one sample. *N*_sig_ is the number of sites with significant changes in editing yield. Points corresponding to such sites are shown with red crosses. (right-panel) A comparison of the number of off-targets induced by delivery of circular adRNAs, linear adRNAs, and linear adRNAs with co-delivered ADAR2 (*1*). All experiments were carried out in HEK293FT cells. (**b**) Engineered cadRNA designs for reducing bystander editing, and associated heatmaps of percent editing within a 60bp window around the target adenosine in the GAPDH and RAB7A transcripts. The positions of adenosines relative to the target adenosine (0) are listed below the heatmap. The strand in pink represents the antisense domain while the target mRNA is in blue. The target adenosine is highlighted in green while the mismatch opposite it is in red. Design 1: Unmodified circular.200.100 antisense. Design 2: Antisense bulges created by positioning guanosines opposite bystander adenosines. Design 3: Loops of size 8bp created at position -6 and +30 relative to the target adenosine. Values represent mean % editing (n=3 for GAPDH and n=2 for RAB7A). All experiments were carried out in HEK293FT cells. (**c**) Plasmid delivered *in situ* cadRNA generation: RNA editing efficiencies across various transcripts observed in HEK293FT and K562 cells via plasmid delivered circular adRNA is shown. Values represent mean +/- SEM (n=3). Associated changes in expression levels of target transcripts as compared to levels seen in untransfected controls is also shown (p=0.2599, p=0.0135, p=0.1982, p=0.7871, p=5145; unpaired t-test, two-tailed). (**d**) *In vitro* transcribed (IVT) circular adRNA generation: Linear forms of twister ribozyme flanked circular adRNAs were transcribed *in vitro* using a T7 polymerase, purified using LiCl and transfected into cells, where they circularize *in situ* by the endogenous RNA ligase RtcB. RNA editing efficiencies across various transcripts observed in HEK293FT and K562 cells via IVT circular adRNA is shown. Values represent mean +/- SEM (n=3). Associated levels of IVT and plasmid delivered circular.200.100 adRNA targeting RAB7A measured in transfected HEK293FT cells is also shown. Values represent mean +/- SEM (n=3).

Next, we confirmed the robustness and generalizability of the cadRNA format via their ability of to successfully edit adenosines in the coding sequence (CDS) of three additional transcripts – GAPDH, ALDOA and DAXX in HEK293FT cells (**Figure 2c**). Furthermore, in addition to delivery via a genetically encoded format in plasmids, we also explored if *in vitro* transcribed (IVT) circular adRNA would be similarly functional. Towards this, IVT cadRNAs were engineered by delivering to cells the twister ribozyme flanked adRNAs in a linear form which then underwent *in situ* circularization in the cells (**Figure 2c**). 24hrs post transfection, we observed robust editing of the RAB7A and GAPDH transcripts using IVT adRNAs in HEK293FTs (**Figure 2d**) and also confirmed circularization of the IVT adRNAs via qPCR. Additionally, the plasmid and IVT adRNAs based editing of RAB7A in K562 cells using electroporation was similarly robust at 90% and 70% RNA editing yields respectively (**Figures 2c, 2d**). Additionally, for a majority of the tested loci we did not observe significant knockdown of the targeted transcripts (**Figure 2c**).

Given the vastly improved efficiency and durability of RNA editing via cadRNAs, we next wondered if these could enable *in vivo* RNA editing. Since no co-delivery of proteins is required, successful demonstration here could enable a powerful gene therapy approach. Additionally, one could leverage for cadRNAs the already established delivery modalities and accruing knowledge from the field of shRNAs and ASOs that similarly only require delivery of nucleic acids to target tissues. To explore this, we first targeted an adenosine in the 3’ UTR of the mPCSK9 transcript via AAV8 mediated delivery of adRNAs to the mouse liver. We systematically compared RNA editing yields via linear.U6+27.100.50, one copy of circular.200.100, and two copies of circular.200.100 guide RNAs (**Figure 3a**). 2 weeks post injections, we harvested mice livers and did not detect any editing in the PBS injected mice, and notably in the mice injected with linear.U6+27.100.50 guide RNAs too we did not measure detectable RNA editing (**Figure 3b**). Excitingly, we observed highly efficient 11% and 38% on-target editing via the single copy and two copy circular.200.100 guide RNAs respectively. We confirmed via qPCR robust expression of the cadRNAs, and noted that addition of a second copy of the circular.200.100 led to a further 3-fold increase in expression levels, together suggesting that persistent and robust guide RNA expression was key to enabling efficient RNA editing (**Figure 3c**). Importantly, we also confirmed that cadRNAs delivered via AAVs did not alter the expression levels of the mPCSK9 transcript in mice livers (**Figure 3d**).

**Figure 3:**
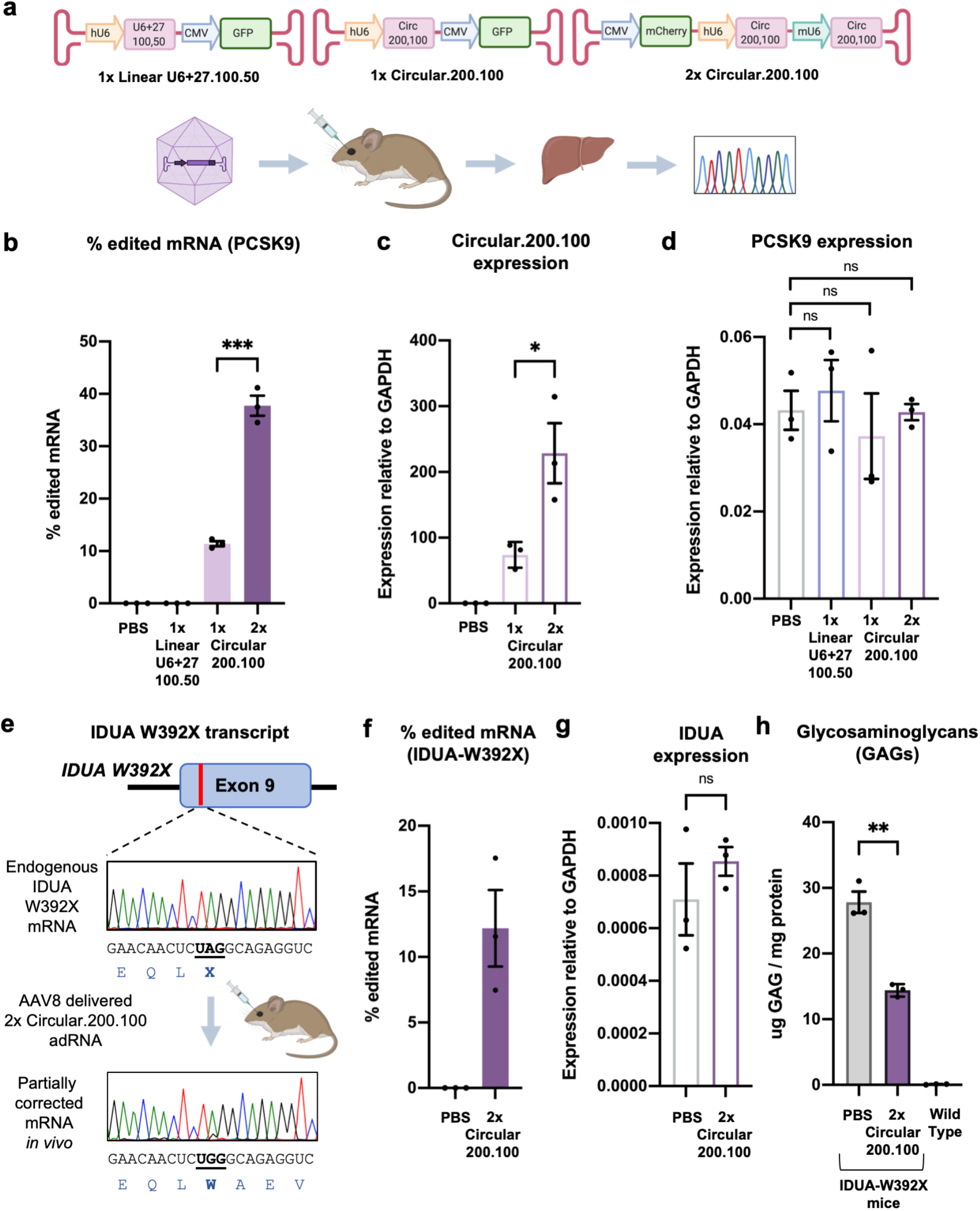
*In vivo* efficacy of cadRNAs. (**a**) (i) AAV vectors used for adRNA delivery. (ii) Schematic of the *in vivo* experiment. (**b**) *In vivo* RNA editing efficiencies of the mPCSK9 transcript in mice livers via systemic delivery of linear and circular adRNAs packaged in AAV8. Values represent mean +/- SEM (n=3; p=0.0002; unpaired t-test, two-tailed). (**c**) Relative expression levels of circular adRNAs. Values represent mean +/- SEM (n=3; p=0.0305; unpaired t-test, two-tailed). (**d**) mPCSK9 transcript levels relative to GAPDH. Values represent mean +/- SEM (n=3; p=0.6179, p=0.6125, p=0.9323; unpaired t-test, two-tailed). (**e**) Schematic of the IDUA-W392X mRNA, and RNA editing experiment. (**f**) *In vivo* UAG-to-UGG RNA editing efficiencies of the IDUA transcript in mice livers via systemic delivery of linear and circular adRNAs packaged in AAV8. Values represent mean +/- SEM (n=3). (**g**) IDUA transcript levels relative to GAPDH. Values represent mean +/- SEM (n=3; p=0.3815; unpaired t-test, two-tailed). (**h**) GAG content in mice livers of PBS injected and AAV8-circular.200.100 injected IDUA-W392X mice. Wild type C57BL/6J mice were included as controls. Values represent mean +/- SEM (n=3; p=0.0014; unpaired t-test, two-tailed).

Building on these results, we next targeted a mouse model of Hurler syndrome. Hurler syndrome is a form of mucopolysaccharidosis type 1 (MPS1), a rare genetic disorder which results in the buildup of large sugar molecules called glycosaminoglycans (GAGs) in lysosomes. This occurs due to a lack of the enzyme alpha-L-iduronidase which is encoded by the IDUA gene. W402X is a commonly occurring mutation in the IDUA gene in Hurler syndrome patients and there exists a corresponding mouse model bearing the IDUA-W392X mutation (*34*) (**Figure 3e**). With a goal to repair the IDUA-W392X premature stop codon, we packaged 2 copies of circular.200.100 guide RNA into AAV8 and injected these into IDUA-W392X mice systemically. Two weeks post injection, we harvested mice livers and observed robust 7-17% correction of the premature stop codon in the mice (**Figure 3f**). Further, we confirmed that expression of the circular.200.100 adRNA did not alter the expression levels of the IDUA transcript (**Figure 3g**). We also measured GAG levels in these mice, and observed in the treated animals these resulted in about 50% less accumulation over the 2 week period than PBS injected mice, indicating successful partial restoration of alpha-L-iduronidase activity (**Figure 3h**).

## DISCUSSION

Use of endogenous ADARs for correction of G-to-A point mutations and premature stop codons carries immense therapeutic potential. However, the relatively short half-life of the guide RNAs limits efficacy. In this study, we engineered circular guide RNAs for recruitment of endogenous ADARs that vastly improve the efficiency and durability of programmable RNA editing. This method is highly specific at the transcriptome-level, and engineering of bulge loops in the antisense domain also enabled high specificity at the transcript-level with significantly reduced bystander adenosine editing. Via AAV-delivered cadRNAs we also demonstrated for the first time robust *in vivo* RNA editing via endogenous ADAR recruitment, including in the IDUA-W392X mouse model of mucopolysaccharidosis type I-Hurler (MPS I-H) syndrome. While cadRNAs provide an exciting format for RNA editing, there are several areas that merit further investigation. Specifically: 1) Improving cadRNA editing yields via coupling to additional ADAR recruitment domains; 2) Enhancing the ability of the antisense region to hybridize to target RNAs via structural pre-straining of the cadRNAs; 3) Assaying the innate immune response to circular RNAs, generated both via genetically encoded vs. IVT formats, and those delivered via viral vs. non-viral modalities (*35*). In particular, for the IVT formats, we anticipate introduction of modified RNA bases might be critical for enhancing adRNA efficacy; 4) Reducing bystander editing by cadRNAs while maintaining on-target editing. This could entail combining both generic as well as target specific approaches, for instance, integrating the loop designs and A-specific bulges; and 5) Monitoring undesired RNAi effects. As noted both in this and our prior work (*1*), while a majority of targets maintained expression levels, for some targets clear RNAi effects were observed via both long-antisense adRNAs and cadRNAs, and correspondingly, modifying those guide designs will be critical to enable efficacious editing. Moving beyond, we anticipate circularization of guide RNAs might also have utility in other transcriptome and genome engineering modalities, such as RNAi, ASOs, and guide RNAs in CRISPR-Cas. In summary, as cadRNAs do not require the need for co-delivery of any effector proteins, and as a targeting moiety also have enhanced persistence in cells, these have the potential for broad utility in programmable RNA editing mediated transient protein modulation, and correction of G-to-A point mutations and premature stop codons for therapeutic applications.

## ACKNOWLEDGEMENTS

We thank Genghao Chen, Lauren Hodge and other members of the Mali lab for discussions, advice and help with experiments. This work was generously supported by UCSD Institutional Funds and NIH grants (R01HG009285, RO1CA222826, RO1GM123313, 1K01DK119687). This publication includes data generated at the UC San Diego IGM Genomics Center utilizing an Illumina NovaSeq 6000 that was purchased with funding from a National Institutes of Health SIG grant (#S10 OD026929). Schematics were created using BioRender.

## AUTHOR CONTRIBUTIONS

D.K. and P.M. conceived the study and wrote the paper. D.K., P.M., J.Y., Y.X., A.S. and Y.S. performed experiments. D.M. quantified RNA-editing activity from RNA-seq data.

## METHODS

### Transfections

Unless otherwise stated, experiments were carried out in HEK293FT cells which were grown in DMEM supplemented with 10% FBS and 1% Antibiotic-Antimycotic (Thermo Fisher) in an incubator at 37 °C and 5% CO_2_ atmosphere. HEK293FT cells were seeded in 24 well plates and transfected using 1000 ng adRNA plasmid or 48 pmol of IVT RNA and 2ul of commercial transfection reagent Lipofectamine 2000 (Thermo Fisher). Cells were transfected at 25-30% confluence. Plasmid transfection experiments were harvested 48 hours post transfections while IVT RNA experiments were harvested 24 hours post transfections. For 96 hour long experiments, cells were passaged at a 1:4 ratio, 48 hours post transfections. Cells after plasmid electroporation were harvested at 48 hours, while IVT RNA experiments were harvested 24 hours post electroporation.

### Electroporation

K562 cells were grown in RPMI supplemented with 10% FBS and 1% Antibiotic-Antimycotic (Thermo Fisher) in an incubator at 37 °C and 5% CO_2_ atmosphere. 200,000 cells were electroporated with 1000 ng adRNA plasmid or 48 pmol of IVT RNA using the Amaxa SF cell Line 4D-Nucleofector X kit (Lonza) as per the manufacturer’s instructions.

### *In vitro* transcription

Sense RNA fragments and and circular adRNA were made by *in vitro* transcription using the HiScribe T7 Quick High Yield RNA Synthesis Kit (NEB) as per the manufacturer’s protocol. DNA templates for the IVT reaction carried the T7 promoter sequence at the 5’ end and were created by PCR amplification of the desired sequence from plasmids or cDNA. PCR products were purified using a PCR Purification Kit (Qiagen) and then used for IVT.

### Luciferase assay

HEK293FT cells were grown in DMEM supplemented with 10% FBS and 1% Antibiotic-Antimycotic (Thermo Fisher) in an incubator at 37 °C and 5% CO_2_ atmosphere. All *in vitro* luciferase experiments were carried out in HEK293FT cells seeded in 96 well plates, at 25-30% confluency, using 200 ng total plasmid and 0.4 μl of commercial transfection reagent Lipofectamine 2000 (Thermo Fisher). Specifically, every well received 100 ng each of the Cluc-W85X(TAG) reporter and the adRNA plasmids. At the same time, every well also received 25 pmol siRNA. 48 hours post transfections, 20 μl of supernatant from cells was added to a Costar black 96 well plate (Corning). For the readout, 50 μl of Cypridina Glow Assay buffer was mixed with 0.5 μl Vargulin substrate (Thermo Fisher) and added to the 96 well plate in the dark. The luminescence was read within 10 minutes on Spectramax i3x or iD3 plate readers (Molecular Devices) with the following settings: 5 s mix before read, 5 s integration time, 1 mm read height.

### Production of AAV vectors

AAV8 particles were produced using HEK293FT cells via the triple-transfection method and purified via an iodixanol gradient. Confluency at transfection was about 50%. Two hours before transfection, cell medium was exchanged with Dulbecco’s modified Eagle’s medium supplemented with 10% fetal bovine serum and 100X Antibiotic-Antimycotic (Gibco). All viruses were produced in 5×15 cm plates, where each plate was transfected with 10 μg of pXR-8, 10 μg of recombinant transfer vector and 10 μg of pHelper vector using polyethylenimine (PEI) (1 μg/μl linear PEI in ultrapure water, pH 7, using hydrochloric acid) at a PEI:DNA mass ratio of 4:1. The mixture was incubated for 10 minutes at room temperature and subsequently applied dropwise onto the cell media. The virus was harvested after 72 hours and purified using an iodixanol density gradient ultracentrifugation method. The virus was then dialyzed with 1× phosphate buffered saline (pH 7.2) supplemented with 50 mM sodium chloride and 0.0001% Pluronic F68 (Thermo Fisher) using 50 kDA filters (Millipore), to a final volume of ∼1 ml, and quantified by quantitative PCR using primers specific to the ITR region, against a standard (ATCC VR-1616): AAV-ITR-F, 5′-CGGCCTCAGTGAGCGA-3′; AAV-ITR-R, 5′-GGAACCCCTAGTGATGGAGTT-3′.

### Animal experiments

All animal procedures were performed in accordance with protocols approved by the Institutional Animal Care and Use Committee of the University of California, San Diego. All mice were acquired from Jackson Labs. AAVs were injected retro-orbitally into both C57BL/6J mice and IDUA-W392X mice (B6.129S-Iduatm1.1Kmke/J), 6-8 weeks of age, at a dose of 1.0E13 vector genomes per mouse. Mice were monitored three times a week for the duration of the experiment (2 weeks).

### GAG assay

The GAG assay was performed following the protocol described in (*36*). Briefly, harvested mouse tissues were homogenized in 1 ml PBS with a syringe and 16 gauge (1.6 mm) needle. Tissue homogenates were then incubated on ice for 20 min with Triton X-100 added to a final concentration of 1%. Protein concentration in the supernatant clarified via centrifugation was estimated using the Bradford assay. Supernatants were digested in 1 mg/ml Proteinase K (Qiagen) for 12 h at 55 °C then boiled for 10 min to inactivate the enzyme. Nucleic acids were digested using Benzonase nuclease (Sigma) at 37 °C for 1 h followed by 10 min boiling to inactivate the enzyme. Total amount of GAG in each sample was measured using the Blyscan GAG assay kit (Biocolor).

### RNA extraction and quantification of editing

RNA from cells was extracted using the RNeasy Mini Kit (Qiagen) while extraction from tissues was carried out using QIAzol Lysis Reagent and purified using RNeasy Plus Universal Mini Kit (Qiagen), according to the manufacturer’s protocol. 500-1000 ng RNA was incubated with 1 μl of 5 μM of a target specific sense RNA (synthesized via IVT) at 95 °C for 3 minutes followed by 4 °C for 5 minutes. This step was carried out to capture the circular adRNA which if tightly bound to the target mRNA would block reverse transcription. cDNA was then synthesized using the Protoscript II First Strand cDNA synthesis Kit (NEB). 1 μl of cDNA was amplified by PCR with primers that amplify about 300-600 bp surrounding the sites of interest (outside the length of the antisense domain) using OneTaq PCR Mix (NEB). The numbers of cycles were tested to ensure that they fell within the linear phase of amplification. PCR products were purified using a PCR Purification Kit (Qiagen) and sent out for Sanger sequencing. The RNA editing efficiency was quantified using the ratio of peak heights G/(A+G). RNA-seq libraries were prepared from 250 ng of RNA, using the NEBNext Poly(A) mRNA magnetic isolation module and NEBNext Ultra II Directional RNA Library Prep Kit for Illumina. Samples were pooled and loaded on an Illumina Novaseq 6000 (100 bp paired-end run) to obtain 40-45 million reads per sample.

### Mapping of RNA-seq reads

Sequence read pairs from stranded RNA-seq libraries were mapped to the reference human genome hg38 by running STAR aligner version 2.7.3a (*37*) with the following command line options: --clip3pAdapterSeq AGATCGGAAGAGCACACGTCTGAACTCCAGTCA AGATCGGAAGAGCGTCGTGTAGGGAAAGAGTGT (to trim Illumina adapter sequences from the 3′ ends of the reads in each pair), --quantMode GeneCounts (to collect read counts for each gene), --alignSJDBoverhangMin 1 (following ENCODE standard practice), --peOverlapNbasesMin=10 --peOverlapMMp=0.05 (to correctly align pairs of overlapping reads), --outSAMmultNmax 1 (to limit output of multimapping reads), --alignEndsType EndToEnd (to avoid soft-clipping of reads), --outFilterMismatchNmax -1 --outFilterMismatchNoverReadLmax 0.2 -- outFilterMultimapNmax 1 (to increase the likelihood of successful alignment for reads containing A-to-I editing events). The genome index for STAR aligner was built using transcript annotations from Gencode release 32 (*38*) Each aligned read was retained for downstream analysis even when the corresponding mate in the pair could not be successfully aligned. Samtools version 1.10 (*39*) was used to sort the aligned reads by genomic coordinate and to mark duplicated single or paired reads. The file ReadsPerGene.out.tab generated by STAR aligner contains three types of read counts for each gene: counts collected without considering read strands, counts based on the first strand of each read pair, and counts based on the second strand. The counts based on the first strand were found to be zero for most genes, while the counts based on the second strand were comparable to the unstranded counts, thus confirming that the sequence of first (second) read in each pair of the stranded RNA-seq libraries had the same orientation as the first (second) cDNA strand, as expected from the NEBNext Ultra II Directional RNA Library Prep Kit.

### Quantification of changes in RNA editing

To quantify significant changes in RNA editing, the BAM files containing reads aligned to the reference genome were processed as follows. Reads marked as duplicates were ignored. To minimize the bias of library size on statistical comparisons between different samples, the remaining reads from each sample were down-sampled, using samtools view with option -s, to the smallest number of such reads available for any sample. The down-sampling fraction used for each sample was calculated by dividing the smallest number of uniquely aligned reads among all samples by the number of uniquely aligned reads available for the sample being down-sampled. However, reads for the control sample, which was used for all comparisons, were not down-sampled.

The first step to quantify A-to-I editing events is to count the actual bases occurring on RNA transcripts at positions that, according to the reference genome, are expected to harbor an adenine base. Thus, for transcripts oriented as the forward (reverse) reference strand, base counts must be collected at reference A-sites (T-sites). As noted above, the first (second) read in each pair of the stranded RNA-seq libraries has the same orientation as the first (second) cDNA strand, i.e., the opposite (same) orientation as the transcript from which each cDNA molecule is synthesized. Also, the Illumina sequencing technology yields a pair of reads from opposite strands of the sequenced DNA molecule. Therefore, to handle transcripts oriented as the forward reference strand, base counts were collected at reference A-sites using the second (first) read in a pair, if that read was mapped to the forward (reverse) reference strand. Conversely, to handle transcripts oriented as the reverse reference strand, base counts were collected at reference T-sites using the first (second) read in a pair, if that read was mapped to the forward (reverse) reference strand.

The C library htslib (github.com/samtools/htslib) was used to enumerate the aligned reads that overlapped each base position in the reference genome. Reference sites covered by less than ten reads were ignored. The value of the SAM tag MD, “String for mismatching positions”, recorded by STAR aligner in each alignment record, was used to determine the reference base at each position of an aligned sequence read. Base deletions and insertions relative to the reference genome were ignored. Sequenced bases with a Phred quality score less than 13 were ignored. For each sample, an initial list of base counts from reads overlapping each selected reference A- and T-site was generated.

The initial lists of base counts from all samples were then used to generate a final list of reference A- and T-sites where such base counts were available for all samples, and where at least one sample had a non-zero count of G (C) at reference A-sites (T-sites).

At each selected reference site in the final list, a pairwise comparison between the base counts for each treatment sample and those for the control sample was carried out using Fisher’s exact test, as implemented in R function fisher.test, with a 2-by-2 contingency table containing the counts of G (C) at reference A-sites (T-sites) in the first row, the counts of all other bases at those sites in the second row, the base counts for the control sample in the first column, and the base counts for the compared treatment sample in the second column. The resulting p-values were adjusted for multiple comparisons using the method of Benjamini and Hochberg (*40*), as implemented in R function p.adjust. The proportion of the number of G (C) bases relative to the number all bases was also calculated at each A-site (T-site). Reference A-sites (T-sites) with a significant change in such base proportion for at least one comparison between a treatment sample and the control sample were selected by requiring an adjusted p-value less than 0.01 and a fold change greater than 1.1 in either direction. To visually compare each treatment sample with the control sample, 2D histograms of the observed base proportions at all reference A- and T-sites in the final list were generated using ggplot2 (*41*). The highlighted point in these histograms corresponds to the base proportions calculated, as described above, for the reference A-site at chr3:128814202 (1-based hg38 coordinate).

